# Potent universal-coronavirus therapeutic activity mediated by direct respiratory administration of a Spike S2 domain-specific human neutralizing monoclonal antibody

**DOI:** 10.1101/2022.03.05.483133

**Authors:** Michael S. Piepenbrink, Jun-Gyu Park, Ashlesha Desphande, Andreas Loos, Chengjin Ye, Madhubanti Basu, Sanghita Sarkar, David Chauvin, Jennifer Woo, Philip Lovalenti, Nathaniel B. Erdmann, Paul A. Goepfert, Vu L. Truong, Richard A. Bowen, Mark R. Walter, Luis Martinez-Sobrido, James J. Kobie

## Abstract

Severe Acute Respiratory Syndrome Coronavirus-2 (SARS-CoV-2) marks the third novel β-coronavirus to cause significant human mortality in the last two decades. Although vaccines are available, too few have been administered worldwide to keep the virus in check and to prevent mutations leading to immune escape. To determine if antibodies could be identified with universal coronavirus activity, plasma from convalescent subjects was screened for IgG against a stabilized pre-fusion SARS-CoV-2 spike S2 domain, which is highly conserved between human β-coronavirus. From these subjects, several S2-specific human monoclonal antibodies (hmAbs) were developed that neutralized SARS-CoV-2 with recognition of all variants of concern (VoC) tested (Beta, Gamma, Delta, Epsilon, and Omicron). The hmAb 1249A8 emerged as the most potent and broad hmAb, able to recognize all human β-coronavirus and neutralize SARS-CoV and MERS-CoV. 1249A8 demonstrated significant prophylactic activity in K18 hACE2 mice infected with SARS-CoV-2 lineage A and lineage B Beta, and Omicron VoC. 1249A8 delivered as a single 4 mg/kg intranasal (i.n.) dose to hamsters 12 hours following infection with SARS-CoV-2 Delta protected them from weight loss, with therapeutic activity further enhanced when combined with 1213H7, an S1-specific neutralizing hmAb. As little as 2 mg/kg of 1249A8 i.n. dose 12 hours following infection with SARS-CoV Urbani strain, protected hamsters from weight loss and significantly reduced upper and lower respiratory viral burden. These results indicate *in vivo* cooperativity between S1 and S2 specific neutralizing hmAbs and that potent universal coronavirus neutralizing mAbs with therapeutic potential can be induced in humans and can guide universal coronavirus vaccine development.

## Introduction

SARS-CoV-2 is the third known emergence of a coronavirus (CoV) with the ability to cause substantial human morbidity and mortality. The SARS-CoV-2 pandemic has resulted in over 5 million deaths worldwide in two years and painfully highlights the vulnerability of humanity to novel CoV. Despite the rapid development of vaccines exhibiting high levels of efficacy and low levels of side effects, global vaccine implementation has been slow. As a result, SARS-CoV-2 virus has infected many human hosts, allowing for the evolution of new variants that have the potential to evade the immune response elicited by previous infection or vaccination.

Although first generation SARS-CoV-2 vaccines have been highly effective at preventing severe disease, including from VoC, the humoral immunity induced by vaccination and natural infection is overwhelmingly dependent on a neutralizing antibody response targeted to the Receptor Binding Domain (RBD) of the Spike (S) glycoprotein. The RBD, which mediates the initial attachment of SARS-CoV-2 to its primary receptor, angiotensin converting enzyme 2 (ACE2), and the S1 domain overall have undergone substantial antigenic diversity since the initial emergence of SARS-CoV-2. Mutations within RBD and S1 dramatically negate the neutralizing activity of plasma antibodies (Abs) against SARS-CoV-2 VoC that are generated from vaccinated or infected individuals and enhance the transmissibility and pathogenicity of VoC (1–4). A good example of significant immune escape is the emergence of the Omicron VoC, with numerous mutations in the RBD that contribute to reduced neutralization by therapeutic human monoclonal antibodies (hmAbs) which are against RBD, and by vaccine and infection-induced plasma Abs (5, 6).

The Spike is assembled as a homotrimer with ~24 molecules located on the surface of each SARS-CoV-2 virion (7). While being synthesized, the S protein is initially cleaved by furin or furin-like proprotein convertase in the Golgi resulting in the S1 domain being non-covalently linked to the S2 domain (i.e. stalk) of the protein (8). Mature viruses are released from infected cells after virus containing vesicles fuse with the cell membrane. CoV S is a class I virus fusion protein (9), with the S1 domain mediating attachment primarily through its RBD, while the S2 domain mediates fusion to the host cell membrane and entry. The SARS-CoV-2 viral fusion mechanism is conserved across numerous pathogen proteins including HIV envelope, influenza hemagglutinin, Ebola GP, and RSV fusion protein (10). While CoV S1 domains exhibit substantial variation to allow recognition of different host receptors, CoV S2 domains are highly conserved overall (90% between SARS-CoV and SARS-CoV-2), which is consistent with their conserved function. For membrane fusion to occur a highly dynamic transformation must occur, requiring precise interactions of various S2 subdomains(11), suggesting there is low tolerance for S2 variation without compromising viral fitness. Antibody responses to S2 are increased in some individuals following SARS-CoV-2 infection and plasma S2 Ab response are associated with survival after coronavirus disease 2019 (COVID-19) infection(12). However, the development of S2-specific Abs following SARS-CoV-2 natural infection or standard vaccination is limited compared to those specific for S1(13, 14). While most S1 Abs are neutralizing, only a small fraction of S2 Abs are neutralizing(13).

Many SARS-CoV-2 neutralizing antibodies also target the N-terminal domain (NTD). While neutralizing Abs to NTE have been identified, new SARS-CoV-2 variants such as Omicron, have gained several mutations in the NTD and RBD of the Spike protein that promote immune evasion. With the substantial number of people unvaccinated and immunocompromised, and emergence of new variants that can evade immune protection, a great need remains for new therapeutics that neutralize SARS-CoV-2, despite its continued evolution. Furthermore, there is a potential treatment advantage of combining Abs that inhibit both ACE2 binding and viral fusion.

Therapeutic mAbs are typically delivered systemically through intravenous infusion or intramuscular injection, which is highly inefficient and slow in achieving optimal concentrations in the respiratory tract for the treatment of respiratory infections(15), including SARS-CoV-2(16). We have previously demonstrated that direct respiratory delivery of a SARS-CoV-2 RBD-specific hmAb enables substantial dose-sparing therapeutic activity in hamsters and here evaluate this delivery mechanism for the treatment of SARS-CoV-2 and SARS-CoV with a S2-specific hmAb.

To more precisely identify the potential of the SARS-CoV-2 S2 specific B cell response, a panel of S2-specific hmAbs were isolated and their molecular features, reactivity profiles, and *in vitro* and *in vivo* antiviral activities were defined. Several of these hmAbs demonstrate broad CoV reactivity, neutralizing, and antibody-dependent phagocytosis activity. With the most potent and broad S2-specific hmAb, 1249A8 exhibiting prophylactic and therapeutic activity against SARS-CoV-2 and SARS-CoV in multiple animal models, demonstrating the universal CoV therapeutic potential of S2 hmAb directly. The combination of 1249A8 with and S1-speciific hmAb 1213H7 was evaluated for possible synergistic effects and delivered to the respiratory tract for improved dose delivery efficiency.

## Results

### Identification and isolation of S2-specific human B cells

To identify SARS-CoV-2 S2-specific human B cells, two complementary recombinant proteins were designed and produced; a pre-fusion state stabilized SARS-CoV-2 (S2-STBL) and a SARS-CoV/SARS-CoV-2 full Spike chimera consisting of SARS-CoV S1 and SARS-CoV-2 S2 (SARS-CoV-1/2 S1S2) (**Figure 1A**). Initial testing of plasma from COVID-19 convalescent patients was performed to identify those with high avidity IgG binding titers against S2 (**Figure 1B**). Using fluorescent S2-STBL and S1S2 chimera tetramers, peripheral blood memory B cells from several subjects were single-cell sorted by flow cytometry (**Figure 1C**) and recombinant fully human IgG1 mAbs (hmAbs) were generated. Seventeen hmAbs with reactivity to SARS-CoV-2 S2 protein resulted (**Figure 1D**). In general, most hmAbs bound commercial preparations of SARS-CoV-2 S, as well as S2-STBL and SARS-CoV-1/2, as shown in the plasma profiling. Binding to S2-STBL and SARS-CoV-1/2 S1S2 was more discriminating, as also evident in the plasma profiling. The previously reported S2-specific hmAb CC40.8 (17) was included as a positive control. Off-target binding to SARS-CoV-2 S1 was not evident.

**Figure 1.**
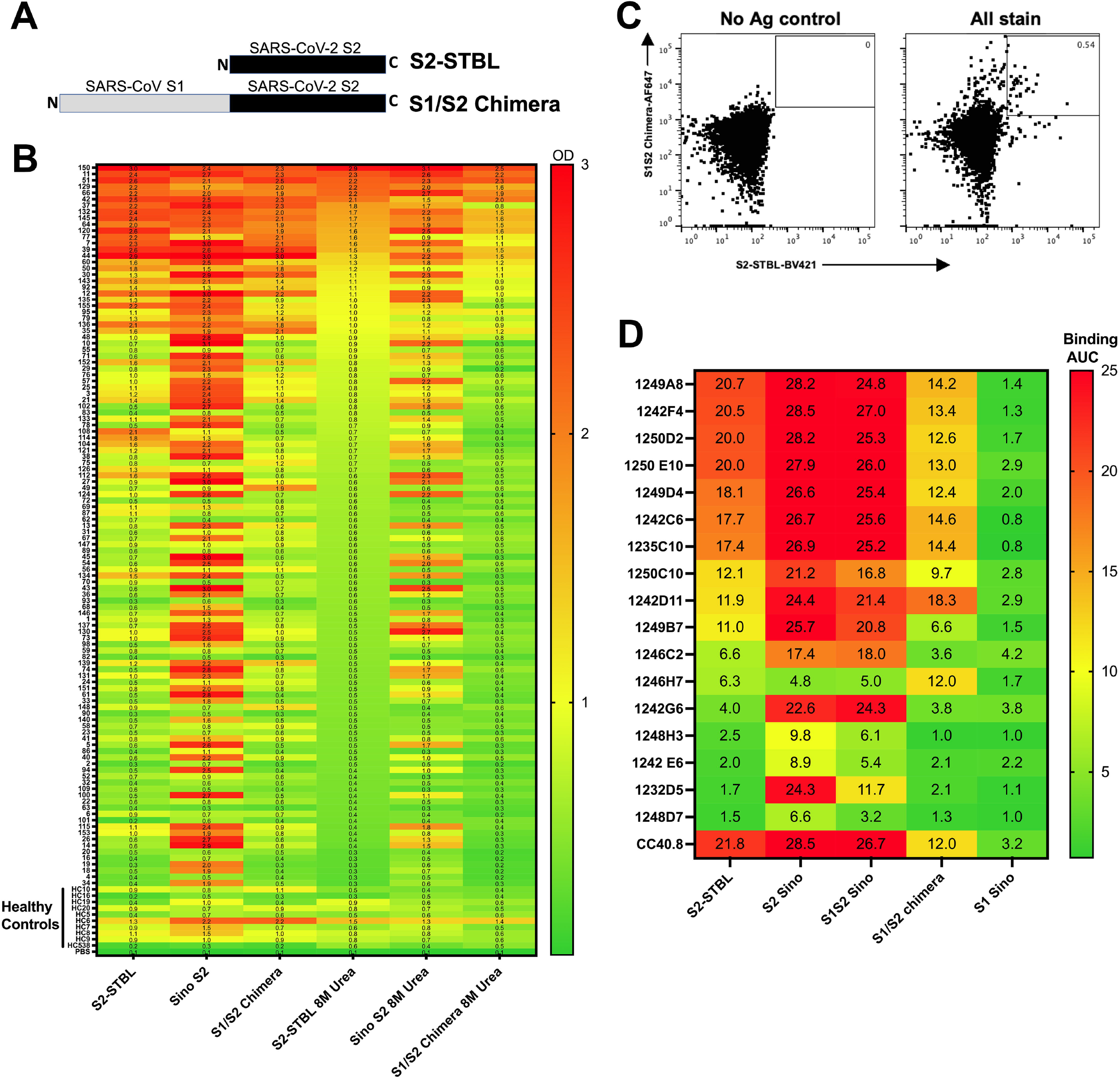
Isolation of SARS-CoV-2 S2-specific human monoclonal antibodies (hmAbs) (**A**) Schematic representation of the S2-STBL and S1/S2 chimera proteins used as baits for ELISA and flow cytometry. **(B)** Human plasma from either convalescent or healthy subjects was diluted 1:1000 in PBS and used in an ELISA against indicated proteins; Absorbance at 450 nM is shown. Each row is an individual subject. **(C)** Representative gating strategy for S2+ B cell isolation. Initial plots are gated on live CD3-CD4-CD14-annexinV-CD19+CD27+ B cells. **(D)** hmAbs were tested at 10 and 1 μg/ml in duplicate by ELISA for binding to indicated protein; area under the curve (AUC) is indicated.

### S2 hmAbs have in vitro SARS-CoV-2 neutralizing and antibody-dependent phagocytosis activity

The functional activity of S2 hmAbs against SARS-CoV-2 was tested. The hmAbs that showed the greatest binding to at least one S2 protein by ELISA were tested by live virus and pseudovirus-based neutralization assays. Several hmAbs did not show neutralization capacity, even at the highest concentration (50 μg/ml) (**Figure 2A+B**). Eight hmAbs demonstrated neutralization of SARS-CoV-2 D614G pseudovirus (PsV) and were tested further, of which four hmAbs (1249A8, 1242C6, 1250D2, and 1235C10) effectively neutralized both PsV and live virus including Beta and Delta VoC. 1249A8 emerged as having the broadest and most potent neutralizing activity, with comparable NT_50_ (neutralization titer at 50% inhibition) to CV3-25, a previously described S2 neutralizing hmAb (18), with the notable exception of SARS-CoV-2 Beta which was not neutralized by CV3-25.

**Figure 2.**
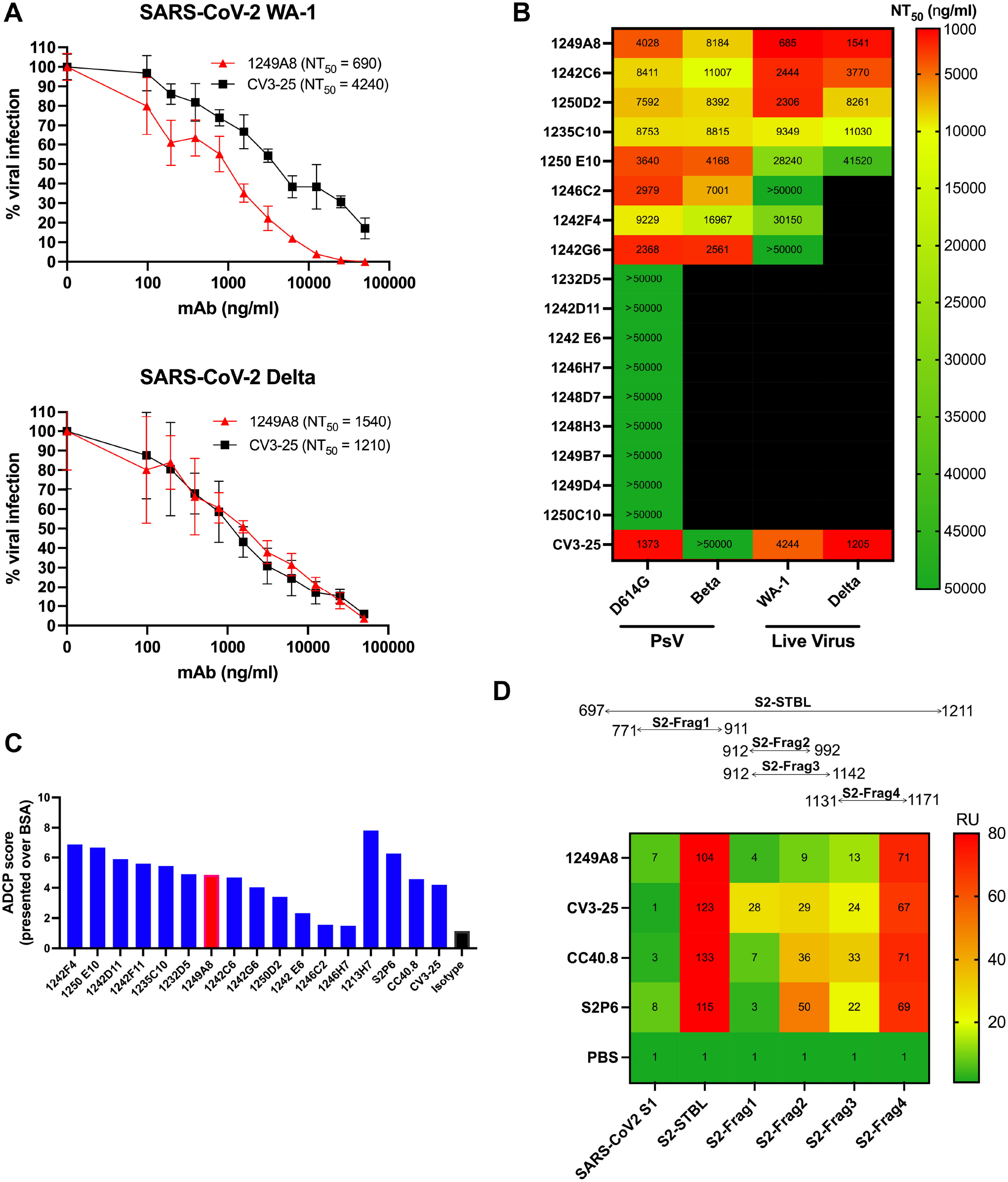
in vitro neutralization and ADCP of SARS-CoV-2 by S2-specific hmAbs. (**A**) SARS-CoV-2 neutralization of S2 hmAbs. Vero E6 cells were infected with SARS-CoV-2 WA-1 or SARS-CoV-2 Delta for 1 h. After 1 h of viral adsorption, the indicated concentrations of S2 hmAbs were added and at 24 h.p.i., infected cells were fixed for virus titration by immunostaining assay. Data was expressed as mean and SD of quadruplicates. (**B**) Summary of viral neutralization (NT_50_) using either pseudovirus representing SARS-CoV-2 D614G mutation or Beta, or live SARS-CoV-2 WA-1 or Delta. (**C**) Ab-dependent cellular phagocytosis (ADCP) assay. SARS-CoV-2 Wuhan-Hu-1 S-coated and BSA-coated beads were incubated with 5 μg/ml hmAb for 2 h and then added to THP-1 cells. After incubation for 3 h at 37°C, cells were assayed for fluorescent bead uptake by flow cytometry. The ADCP score of each mAb was calculated by multiplying the percentage of bead positive cells (frequency of phagocytosis) by the mean fluorescence intensity (MFI) of the beads (degree of phagocytosis) and dividing by 10^6^. (**D**) Binding to S2 protein fragments by hmAbs (5 μg/ml) determined by ELISA.

The Fc effector function of the S2 hmAbs was assessed by antibody dependent cellular phagocytosis (ADCP) of SARS-CoV-2 Wuhan-Hu-1 Spike coated beads (**Figure 2C**). 1242F4 and 1250E10 had the highest ADCP activity, similar to the previously described S2-specific hmAb S2P6 (19). Both 1246C2 and 1246H7 had activity that was only slightly higher than the isotype control indicating very little Fc effector function, and consistent with their limited binding and neutralizing activity. 1249A8 had 4.2 fold greater ADCP activity than the isotype control which was similar to the S2 hmAbs CC40.8 and CV3-25. The SARS-CoV-2 RBD specific mAb 1213H7 (16, 20, 21), had the greatest (~8 times greater than isotype) ADCP activity. Together these results suggest S2 hmAbs have the potential to eliminate SARS-CoV-2 through both neutralization and Fc-dependent effector functions. Four S2 protein fragments (S2 Frag1-Frag4) that cover different regions of the S2 amino acid sequence were produced to identify the binding epitope of 1249A8 (**Figure 2D**). S2 fragment binding assays localized the 1249A8 binding epitope to S2 residues 1131-1171 (S2-Frag4), which contains the conserved stem helix region (residues 1148-1158) of S2, previously reported to be recognized by mAbs CV3-25 (18), CC40.8 (17), and S2P6 (19).

### Molecular characteristics of S2 hmAbs

The most potent neutralizing hmAb, 1249A8 was isolated from an IgG1 expressing B cell and exhibited substantial somatic hypermutation including 16.7% amino acid mutation from germline in the heavy chain variable region, and 7.6% amino acid mutation from germline in the light chain variable region (**Table 1**). The 1249A8 hmAb is member of the same clonal lineage that includes 1242C6, 1250D2, 1242F4, 1249D4, and 1249B7. This shared lineage utilizes VH1-46 heavy chain gene and Vκ3-20 light chain gene, with all members isolated from IgG1 expressing B cells and demonstrating substantial somatic hypermutation in the heavy (15.6-18.8% AA) and light (10.4-13.5% AA) from germline.

**Table 1.**
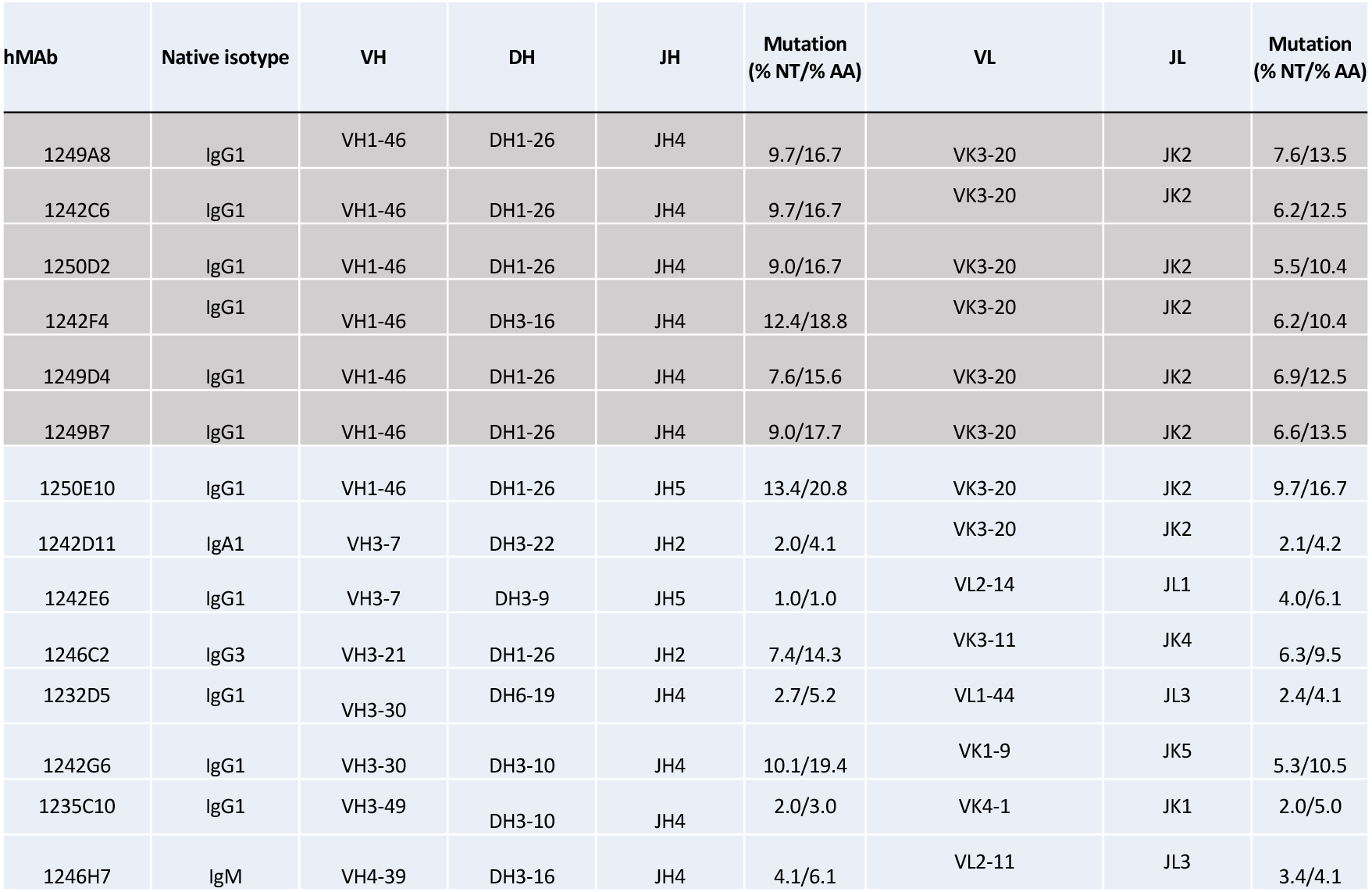
Molecular characteristics of S2 specific hmAbs.

### 1249A8 hmAb protects from SARS-CoV2 infection

The K18 human ACE2 transgenic mouse model was utilized to determine the prophylactic activity of 1249A8 hmAb. Mice were treated with a single dose of 1249A8 intraperitoneally (IP), and 12 hours later challenged with both rSARS-CoV-2 WA-1/Venus and rSARS-CoV-2 Beta/mCherry reporter viruses(21). Mice were also treated alone or in combination with a modest dose of 1213H7 (5 mg/kg), a broad and potent SARS-CoV-2 RBD specific hmAb we have previously described (16, 20, 21). All mice treated with the isotype control hmAb had declining body weight following infection that required euthanasia by D9 (**Figure 3A+B**). Mice treated with 10 mg/kg 1249A8 showed a milder weight loss with 60% of the mice surviving. Mice treated with 40 mg/kg of 1249A8 prior to infection, as well as those treated with 1213H7 or the combination of both did not have weight loss and all survived. Both 10 mg/kg and 40 mg/kg 1249A8 significantly (p<0.05) reduced nasal virus at day (D) 2, with 1249A8 and 1213H7 combination treated mice not having detectable virus, with all 1249A8 treated mice having viral titer below the limit of detection at D4 (**Figure 3C**), with viral burden dominated by rSARS-CoV-2 Beta/mCherry, as previously described(21). Lungs of mice that were treated with the isotype control mAb showed intense fluorescent radiance for both rSARS-CoV-2 WA-1/Venus and rSARS-CoV-2 Beta/mCherry in left and right hemispheres by D2 following infection and markedly increased at D4, and minimally visually evident in the 1249A8, 1213H7, and combination treated mice (**Figure 3D**). Lung viral titer was reduced in mice treated with either 1249A8 doses by ~2 log at D2, and to below detection limit at D4 compared to isotype control hmAb treated mice (**Figure 3E**). The reduction in viral burden by 1249A8 was for both rSARS-CoV2 WA-1/Venus and rSARS-CoV2 Beta/mCherry (**Figure 3E, F, G**) and was consistent with significant (p<0.05) reduction in lung pathology (**Figure 3H**). These results indicate that 1249A8 alone and in combination with the RBD mAb 1213H7 can broadly limit SARS-CoV-2 upper and lower respiratory viral burden and lung pathology, including the original lineage A and the lineage B Beta VoC.

**Figure 3.**
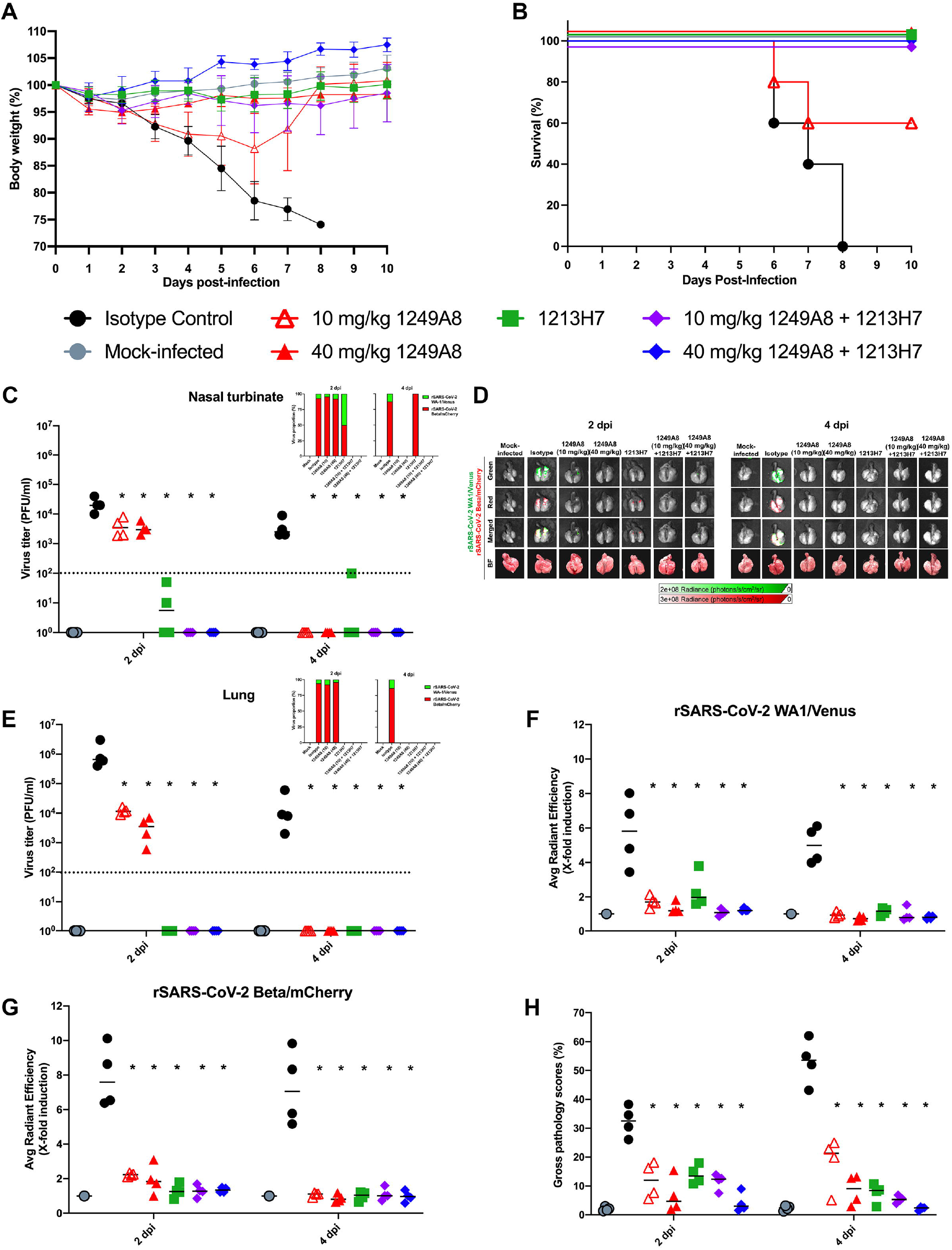
Prophylactic activity of 1249A8 hmAb against rSARS-CoV-2 WA1-Venus and rSARS-CoV-2 Beta-mCherry in K18 ACE2 transgenic mice model. Female K18 hACE2 transgenic mice were treated i.p. with 1249A8 (10 mg/kg or 40 mg/kg), 1213H7 (5 mg/kg), alone or in combination, or isotype control hmAb (40 mg/kg), followed by infection with both rSARS-CoV-2 Venus and rSARS-CoV-2 Beta/mCherry Beta. Body weight (**A**) and survival (**B**) were evaluated at the indicated days post-infection. Mice that loss >25% of their body weight were humanely euthanized. Error bars represent standard deviations (SEM) of the mean for each group of mice. Viral titers in the nasal turbinate (**C**) and lung (**E**) at 2 and 4 DPI were determined by plaque assay in Vero E6 cells. Bars indicate the mean and SD of lung virus titers. Proportion of rSARS-CoV-2 WA-1/Venus and rSARS-CoV-2 Beta/mCherry determined by fluorescence (insets). (**D**) At 2 and 4 DPI, lungs were collected to determine Venus and mCherry fluorescence expression using an Ami HT imaging system. BF, bright field. (**F**) Venus (Green) and (**G**) mCherry (Red) radiance values were quantified based on the mean values for the regions of interest in mouse lungs. Mean values were normalized to the autofluorescence in mock-infected mice at each time point and were used to calculate fold induction. (**H**) Gross pathological scores in the lungs of mock-infected and rSARS-CoV-2-infected K18 hACE2 transgenic mice were calculated based on the percentage of area of the lungs affected by infection. Dotted line indicates limit of detection. * indicates p<0.05 as compared to isotype control hmAb.

### S2 hmAbs have broad β-coronavirus in vitro activity

Given the high conservation of S2 across CoV, the S2 hmAbs were evaluated for their binding and neutralization breadth against diverse SARS-CoV-2 variants and CoV. Vero E6 cells were infected with SARS-CoV-2, SARS-CoV-2 variants, SARS-CoV, and MERS-CoV and binding assessed by immunofluorescence assay (IFA). Six of the hmAbs showed binding to all SARS-CoV-2 isolates, however the hmAb 1242G6 and 1246C2 bound poorly to SARS-CoV-2 infected cells (**Figure 4A**), consistent with their weak neutralizing activity. 1242C6, 1242F4, 1249A8 and 1250D2 all bound to SARS-CoV and MERS-CoV infected cells. CV3-25 had limited binding to SARS-CoV infected cells and no binding to MERS-CoV infected cells.

**Figure 4.**
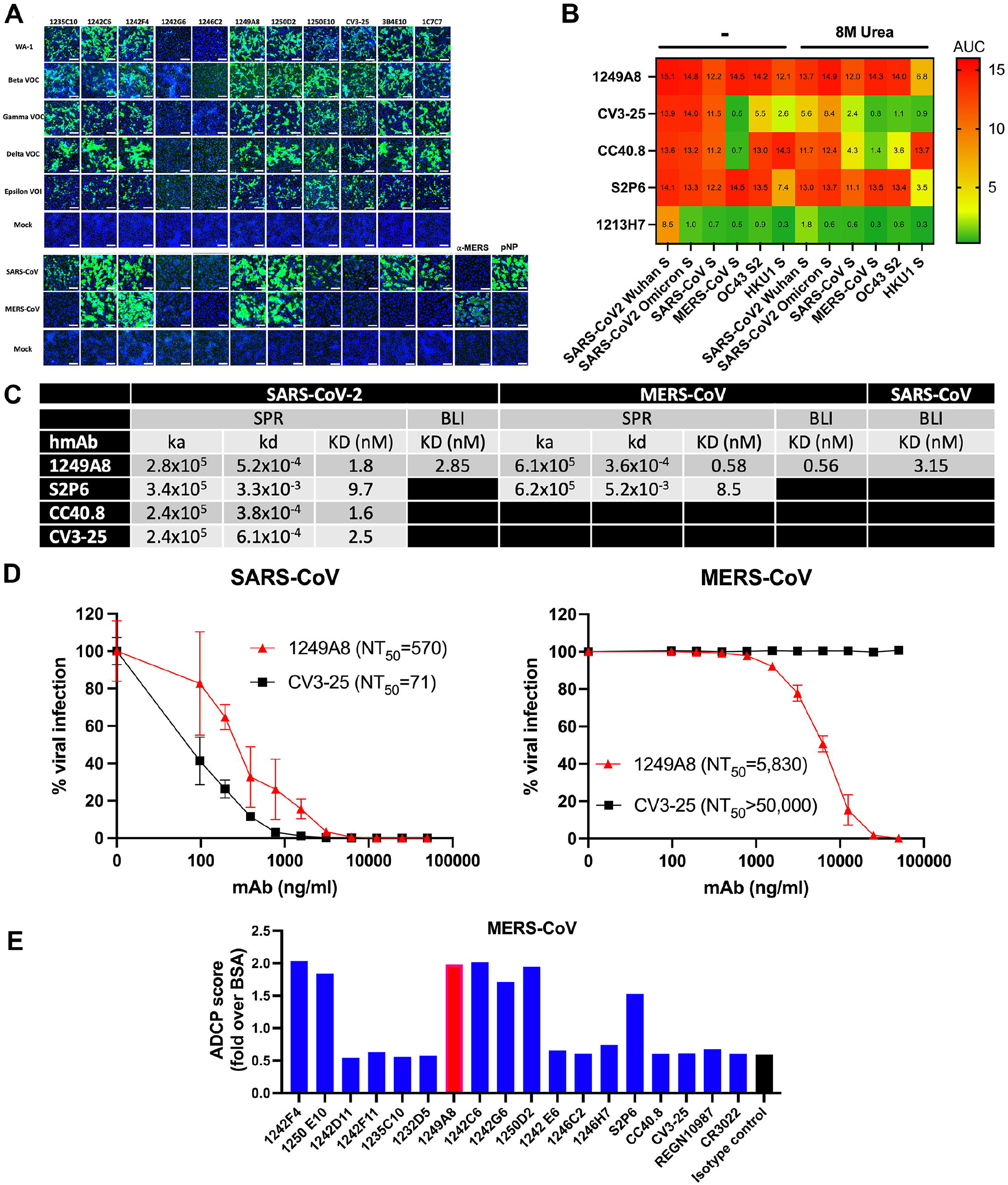
Universal β-coronavirus *in vitro* activity of 1249A8 hmAb. (**A**) Confluent monolayers of Vero E6 cells were infected (MOI 0.1) with SARS-CoV-2 WA-1, Beta (B. 1.351), Gamma (P.1), or Epsilon (B.1.427/B. 1.429). Mock-infected cells (bottom) were included as control. In a separate experiment, confluent monolayers of Vero E6 cells were infected (MOI 0.1) with SARS-CoV (Urbani v2163) or MERS-CoV (recombinant MERS-CoV-RFP delta ORF5 ic). Cells were incubated with the indicated primary S2 hmAb (1 μg/ml) and developed with a FITC-conjugated secondary antihuman Ab. 4,6-Diamidino-2-phenylindole (DAPI; blue) was used for nuclear stain. As internal control, mock and SARS-CoV-2 WA-1 infected cells were stained with the mAb 3B4E10 or with a SARS-CoV cross-reactive NP mAb (1C7C7) or a Ͱ-MERS NP pAb. Scale bars indicate 50 μm. (**B**) Binding of hmAb 1249A8 and other known S2-specific mAb to the Spike proteins of β-coronaviruses as determined by ELISA in the presence or absence of 8M urea. (**C**) Summary table of Surface Plasmon Resonance (SPR) and Biolayer Interferometry (BLI) of 1249A8 against the Spike protein of SARS-CoV-2, MERS-CoV, and SARS-CoV. (**D**) Viral neutralization by 1249A8 and CV3-25. Vero E6 cells were infected with 100 PFU of SARS-CoV (Urbani v2163) or MERS-CoV (recombinant MERS-CoV-RFP delta ORF5 ic) for 1 h. After 1 h of viral adsorption, the indicated concentrations of S2 hmAbs were added. At 24 h.p.i., infected cells were fixed for virus titration by immunostaining assay. (**E**) ADCP of S2-specific mAb against MERS-CoV Spike coated beads.

The breadth of the S2 binding activity of 1249A8 was further evaluated by avidity ELISA in the presence of urea, confirming its binding to SARS-CoV and MERS-CoV Spike, and also demonstrating its binding to OC43 and HKU-1 seasonal β-CoV (**Figure 4B**). No binding to the seasonal α-CoV 229E or NL63 Spike proteins was detected (not shown). 1249A8 uniquely had substantial broad binding to recombinant S proteins from MERS-CoV, OC43, and HKU1 compared to CV3-25 and CC40.8 which did not recognize MERS-CoV S, and S2P6 which had lower reactivity to HKU-1 S compared to 1249A8 (**Figure 4B**). Consistent with the ELISA data, SPR/BLI studies 1249A8 exhibits high affinity for SARS-CoV-2 (*K*D=1.8 nM), SARS-CoV (*K*D=3.2 nM) and MERS-CoV (*K*D=0.58 nM) Spikes. 1249A8 had higher affinity to SARS-CoV-2 and MERS-CoV Spikes as compared to S2P6, as a result of a S2P6 having a faster off-rate (*k*d) (**Figure 4C**). 1249A8 effectively neutralizes both live SARS-CoV (NT_50_=570 ng/ml) and MERS-CoV (NT_50_=5,830 ng/ml) (**Figure 4D**). As expected based on binding activity, CV3-25 did not neutralize MERS-CoV. Additionally, 1249A8 has ADCP activity against MERS-CoV Spike coated beads (**Figure 4E**). These results indicate that several SARS-CoV-2 S2 specific hmAbs have broad beta-CoV reactivity, with 1249A8 demonstrating universal β-CoV functional activity.

### Co-operative S1 and S2 neutralizing mAb in vitro and in vivo activity against SARS-CoV-2 Omicron

The emergence of the SARS-CoV-2 Omicron VoC and its substantial evasion of neutralizing antibodies (2, 5, 6) necessitated testing of 1249A8. 1249A8 retains high affinity (KD = 0.52 nM) for the SARS-CoV-2 Omicron Spike (**Figure 5A**) and neutralizing activity (NT_50_ = 2407 ng/ml) against live SARS-CoV-2 Omicron virus (**Figure 5B**). We also observed potent SARS-CoV-2 Omicron neutralization of 1213H7 (NT_50_ = 64 ng/ml). As clinical development of an S2 mAb would likely include a RBD specific mAb, and as these mAbs target distinct Spike domains (S1 and S2) and steps in the infection process (attachment and fusion) we sought to determine their combinatorial activity. In the presence of 50 ng/ml 1213H7, the NT_50_ of 1249A8 against SARS-CoV-2 Omicron was reduced to 1338 ng/ml, and complementarily, in the presence of 2000 ng/ml of 1249A8 the NT_50_ of 1213H7 was reduced to 26 ng/ml, with similar effect observed for SARS-CoV-2 WA-1 (not shown), suggesting co-operative activity of the S2 and RBD mAbs in neutralizing SARS-CoV-2. Treatment of K18 hACE2 mice with 1249A8 alone i.p. prior to challenge with 10^5^ plaque forming units (PFU) of SARS-CoV-2 Omicron significantly reduced upper and lower respiratory viral burden compared to isotype control treated mice (**Figure 5C+D**). Treatment with 1213H7 alone i.p. also significantly reduced upper and lower respiratory viral burden. The combination of 1249A8 and 1213H7 significantly reduced viral burden, being more pronounced when the hmAbs were administered directly to the respiratory tract through intranasal (i.n.) delivery, with 50% of the mice not having detectable virus in the nasal turbinate at D2 and D4, and lungs at day 2. None of the mice treated i.n. with the 1249A8 and 1213H7 combination had detectable virus in the lungs at D4. This was consistent with the significant reduction in lung pathology in these mice (**Figure 5E**). These results indicate that direct respiratory administration of the RBD and S2 mAb cocktail of 1213H7 and 1249A8, respectively significantly reduce SARS-CoV-2 Omicron viral burden through a cooperative effect.

**Figure 5.**
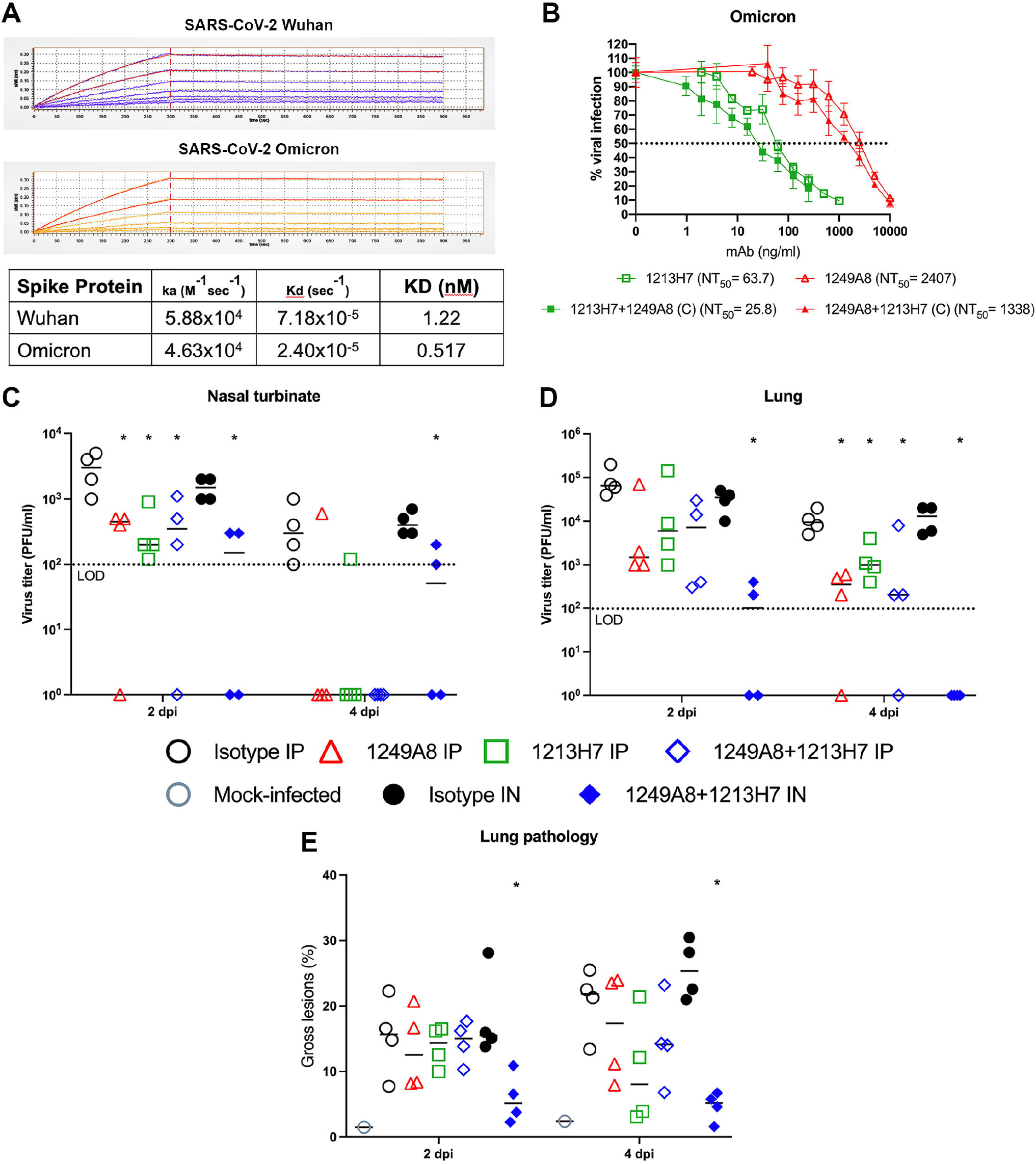
Neutralization and prophylactic *in vivo* activity of 1249A8 and 1213H7 against SARS-CoV-2 Omicron. (**A**) Binding of 1249A8 (60, 40, 27, 18, 12 nM) to SARS-CoV-2 Wuhan and Omicron Spike protein determined by BLI. (**B**) Vero AT cells were infected with 600 pfu SARS-CoV-2 Omicron (BEIR) and after 1 h of viral adsorption, the indicated mAb(s) was added and at 24 h.p.i infected cells were fixed for virus titration by immunostaining assay. 1213H7 and 1249A8 were tested alone (open symbols) and together keeping 1213H7 constant (50 ng/ml) or 1249A8 constant (2 μg/ml) and titrating the reciprocal mAb (closed symbols). Resulting NT_50_ (ng/ml) are indicated. K18 hACE2 mice were treated with 1249A8 (40 mg/kg), 1213H7 (10 mg/kg), or isotype control mAb (40 mg/kg) either alone or in combination i.p. or i.n. as indicated and 24 h later challenged i.n. with 10^5^ PFU SARS-CoV-2 Omicron (BEIR) and virus titer in nasal turbinates (**C**) and lungs (**D**) determined at 2 and 4 dpi by plaque assay and gross lung pathology measured (**E**). Each symbol represents and individual animal. Dotted line indicates limit of detection. *indicates p<0.05 compared to isotype control group as determined by t test.

### Direct respiratory administration of 1249A8 has broad β-coronavirus therapeutic activity

Given the broad β-CoV activity of 1249A8 *in vitro* and its demonstrated prophylactic activity against SARS-CoV-2 WA-1, Beta, and Omicron in K18 hACE2 mice, we evaluated its therapeutic potential in hamsters when delivered directly to the respiratory tract. Hamsters were infected with SARS-CoV-2 Delta and 12 h p.i. were treated with a single dose of hmAb delivered intranasally. Isotype control hmAb treated and untreated hamsters exhibited ~15% body weight loss within 6 days post infection (d p.i.), with minimal weight loss in hamsters treated with the 1249A8 or 1213H7 alone, or in combination (**Figure 6A**). Combined treatment with 8 mg/kg of 1249A8 and 2 mg/kg 1213H7 significantly reduced upper respiratory viral burden by ~3 logs (**Figure 6B**) and lower respiratory viral burden by ~6 logs (**Figure 6C+D**) compared to control groups.

**Figure 6.**
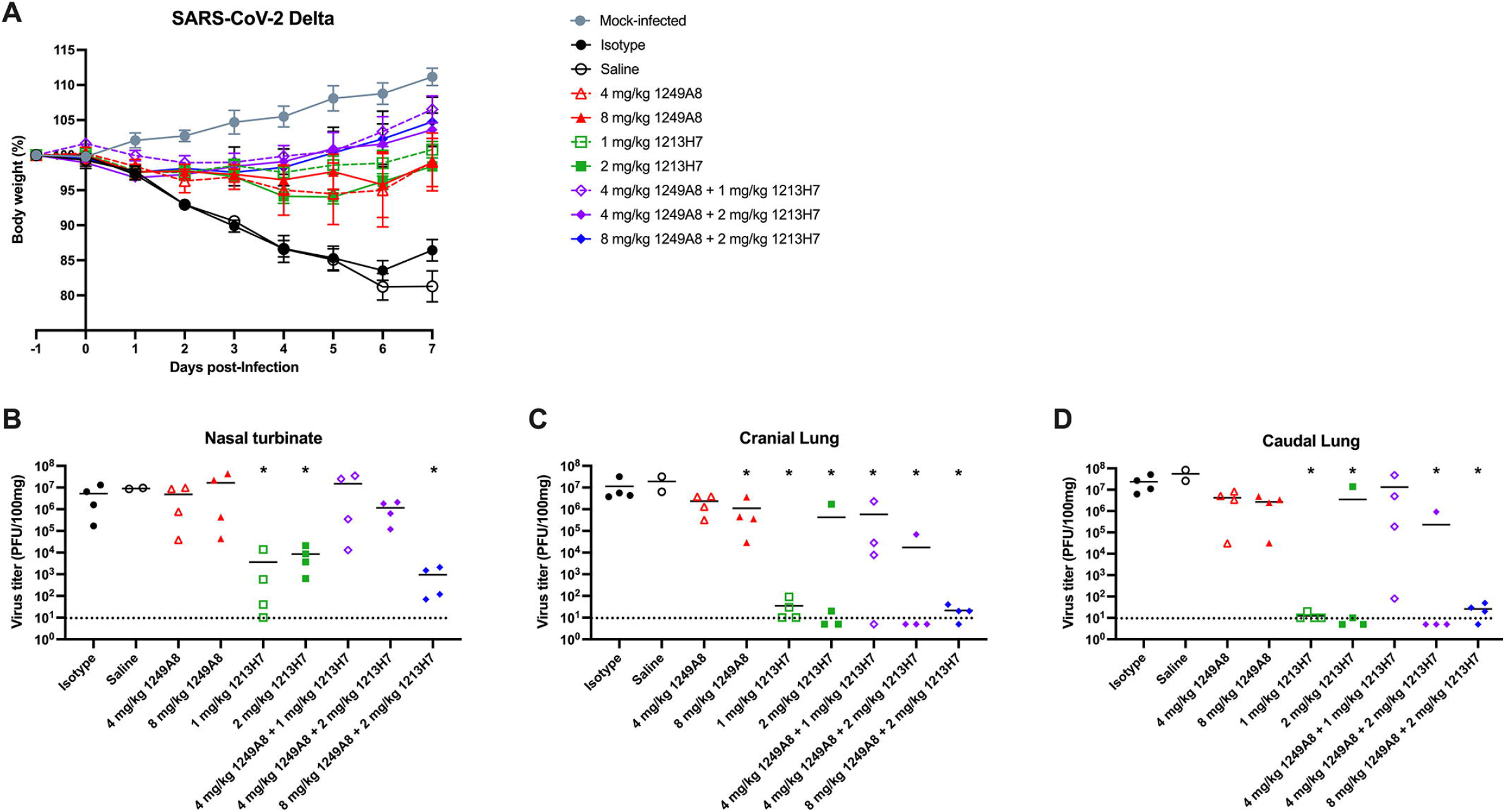
Therapeutic activity of intranasal 1249A8 and 1213H7 in hamsters infected with SARS-CoV-2 Delta. Golden Syrian hamsters were infected i.n. with 10^4^ CCID_50_ SARS-CoV-2 Delta and 12 h p.i. treated i.n. with a single dose of indicated mAb(s). n=4-8 per group. (**A**) Body weight was measured daily. Mean ± SEM indicated. Nasal turbinate (**B**), cranial lung (**C**), and caudal lung (**D**) viral titers were measured at 3 d p.i. by plaque assay. Each symbol represents an individual animal. Dotted line indicates limit of detection. *indicates p<0.05 compared to isotype control group as determined by t test.

To assess pan β-CoV therapeutic activity, hamsters were infected with SARS-CoV, Urbani strain, and 12 h p.i. treated similarly with a single dose of hmAb delivered intranasally. As expected, untreated hamsters and those treated isotype control hmAb lost 15 to 20% of body weight by 7 d p.i.. Hamsters treated with 2, 4 or 8 mg/kg of 1249A8 had <5% weight loss, and those treated with 8 mg/kg 1249A8 alone or in combination with 2 mg/kg of 1213H7 actually gained weight by 7 d p.i. (**Figure 7A**). A significant reduction in upper respiratory viral burden between D1-D3 as determined by oropharyngeal swabbing was evident in all hmAb treated groups (**Figure 7B**). At D3 significant reduction in lung viral titer was evident in hamsters treated with 1249A8 alone and in combination with 1213H7 (**Figure 7DE**). These results indicate the S2 hmAb 1249A8 has broad β-CoV *in vivo* therapeutic activity.

**Figure 7.**
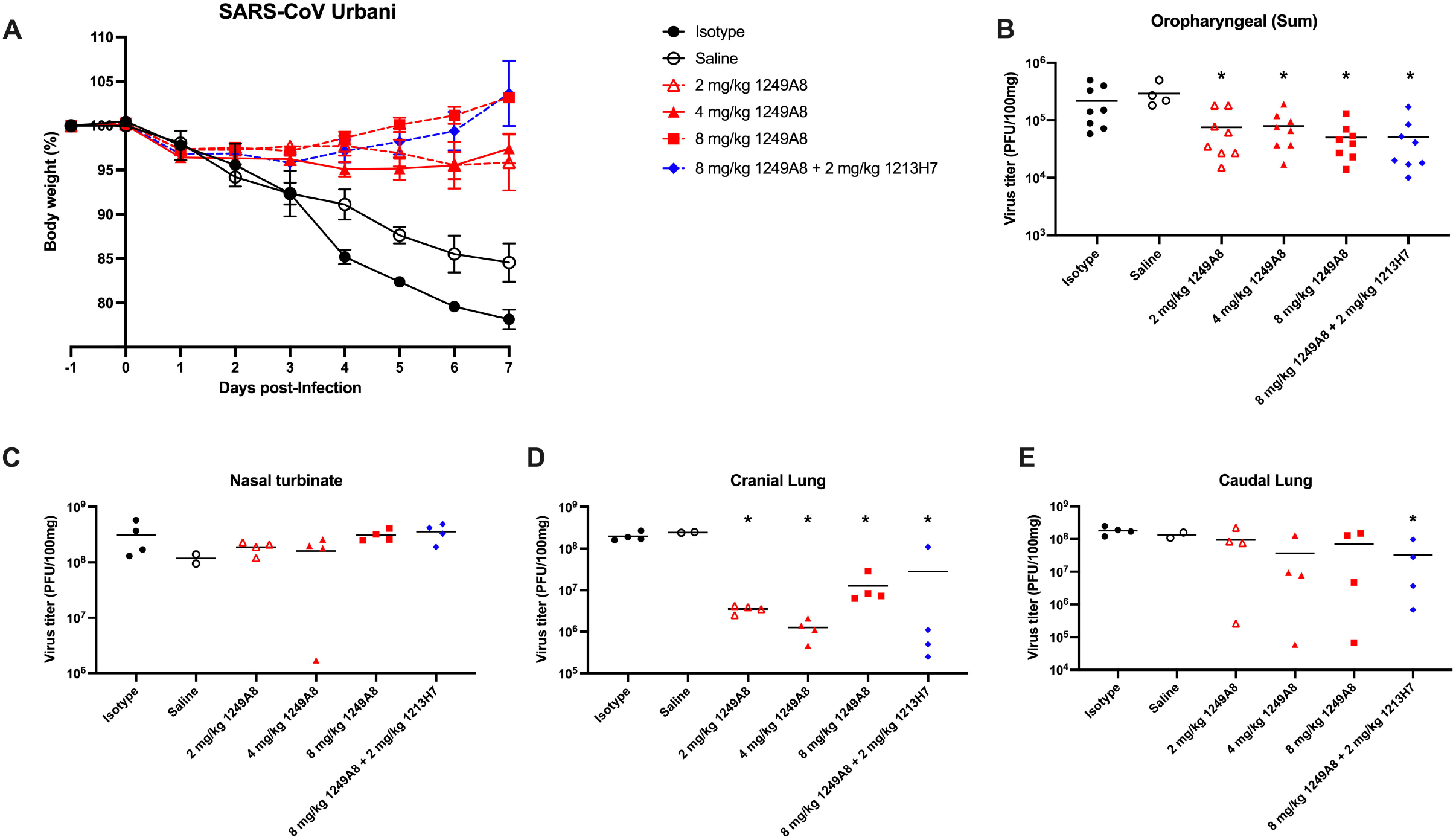
Therapeutic activity of intranasal 1249A8 and 1213H7 in hamsters infected with SARS-CoV. Golden Syrian hamsters were infected i.n. with 10^4^ pfu SARS-CoV (SARS-Urbani) and 12 h p.i. treated i.n. with a single dose of indicated mAb(s). n=4-8 per group. (**A**) Body weight was measured daily. Mean ± SEM indicated. (**B**) Oropharyngeal swabs were collected days 1, 2, and 3 p.i. and sum of virus titer indicated. Nasal turbinate (**C**), cranial lung (**C**), and caudal lung (**E**) viral titers were measured at 3 d p.i. by plaque assay. Each symbol represents an individual animal. *indicates p<0.05 compared to isotype control group as determined by t test

## Discussion

SARS-CoV-2 marks the third time in the last two decades a β-CoV has caused significant mortality in humans. SARS-CoV was originally discovered in Guangdong Province of China in 2002 and spread to five continents but no new cases have been detected since 2004. In 2012 MERS-CoV emerged in the Arabian Peninsula and continues to circulate today. SARS-CoV-2 has caused the most infections and deaths worldwide, new variants pose the risk of evading the immune system even in vaccinated and previously infected individuals, and there remains the potential for other genetically distinct CoV to emerge as new pandemic strains in the future. For these reasons, finding new therapeutic and prophylactic drugs, and vaccine strategies that have universal activity against CoV is essential for protecting humanity.

In SARS-CoV-2 infection, the RBD is immunodominant, having several antigenic sites that account for 90% of the neutralizing activity of convalescent plasma (22). Unfortunately, β-CoV cross-reactive Spike S1/RBD neutralizing antibodies are rare and most S1/RBD mAbs have a very narrow specificity. The Spike protein S2 domain is more highly conserved across β-CoV, and here we have identified several hmAbs from convalescent patients that are highly cross reactive for all variants of SARS-CoV-2 tested. Further, four of these hmAbs also showed cross reactivity to SARS-CoV and MERS-CoV, with 1249A8 hmAb also effectively neutralizing SARS-CoV and MERS-CoV and mitigating SARS-CoV-2 and SARS-CoV infection in animals.

A few S2-specific hmAbs with broad activity have been reported in the literature, and we have included them as possible for comparison in our *in vitro* assessments of 1249A8. 1249A8 is distinct from CV3-25 in its ability to neutralize SARS-CoV-2 Beta VoC, more potent neutralization of SARS-CoV-2 WA-1 consistent with its more potent protection from weight loss and death in K18 hACE2 mice (23), increased binding to OC43 and HKU-1 S, and its ability to bind and neutralize MERS-CoV(24). 1249A8 is distinct from S2P6 in its higher affinity for SARS-CoV-2 S and MERS-CoV S, increased binding to HKU-1 S2, and ability to significantly prevent viral burden from SARS-CoV-2 Beta VoC infection(19). 1249A8 is distinct from CC40.8 in its ability to bind MERS-CoV S(17). Together these findings suggest that 1249A8 is a uniquely broad and potent S2 hmAb that universally recognizes all human β-CoV. 1213H7 demonstrated remarkable breadth for an RBD-specific hmAb in its ability to neutralize all SARS-CoV-2 variants tested including recent Omicron VoC, which evaded many of the hmAbs that were being used clinically for treating COVID-19(25). This suggests that 1213H7 is well suited for the S1 targeting component of an S1/S2 hmAb cocktail for the treatment and prevention of SARS-CoV-2.

Our results suggest that 1249A8 recognizes a yet undefined epitope within stem-helix/HR2 region of S2, and crystallography efforts to precise determine the epitope are ongoing. The ability of 1249A8 to directly neutralize β-CoV infection *in vitro* suggests its binding directly inhibits the post-attachment fusion process, however it remains to be determined at which step(s) such as fusion peptide insertion or 6-helix bundle formation it acts to prevent successful membrane fusion. Finer resolution of the mechanisms of S2 Ab mediated direct inhibition of fusion, and resolution of S2 epitope dynamics in native conformations is needed to adequately identify key β-CoV vulnerabilities.

Although direct inhibition of the viral attachment or fusion is most desired for the antiviral activities of Abs, through Fc fragment engagement, antibodies which may not necessarily bind to neutralizing motifs on infected cells can contribute to overall viral clearance. Antibodies against the S2 portion of the S protein tend to be of lower neutralizing potency than antibodies targeting either the NTD or RBD, so Fc effector function may be of greater importance overall to the S2 Ab response. Although we demonstrated that 1249A8 (as well as others) has ADCP activity against SARS-CoV-2 and MERS-CoV, we do not know to what extent effector function plays in viral clearance. To limit effector function, Ullah et al.(23) introduced the LALA mutation into CV3-25 and found an increase in viral replication, invasion of the brain and body weight loss, while the GASDALIE mutation, which increases Fc effector function, decreased viral dissemination, provided 100% protection, leading to transient body weight loss, decreased viral titers and inflammatory cytokines. Likewise, when S2P6 was expressed as a hamster IgG because human IgG activates Fc receptors poorly in the hamster infection model, Pinto et al.(19) found that the hybrid mAb reduced replicating viral titers to a greater degree than the human counterpart. This context suggests that the Fc-effector function of 1249A8 may contribute to *in vivo* viral clearance and warrants definitive assessment.

The substantial somatic hypermutation that is observed in 1249A8 including 16.7% VH and 13.5% Vk amino acid mutation from germline is higher than would be expected after a primary viral infection. And given its cross-reactivity with seasonal CoV OC43 and HKU-1 suggests that 1249A8 arose from a pre-existing memory B cell that was present in this individual prior to SARS-CoV-2 infection, with this lineage likely originating as a naïve B cell responding to a seasonal β-CoV infection that was expanded further as a consequence of SARS-CoV-2 infection. Given the sequence conservation in S2 among β-CoV, and in light of recent work by Pinto et al., demonstrating S2P6 likely arose from an OC43-specific naïve B cell(19), it remains to be determined if there are constraints in the development of *de novo* S2 memory B cells and Ab responses following SARS-CoV-2 primary infection or vaccination. Determining whether such constraints are a consequence of a lifetime of seasonal β-CoV imprinting or limited immunogenicity of *de novo* SARS-CoV-2 S2 epitopes will be a fundamental for developing universal S2-based CoV vaccines that robustly and reproducibly confer protection in humanities landscape of variable CoV pre-existing immunity.

CoV utilize a variety of host receptors to gain entry, infect, and disseminate. The seasonal CoV including β-CoV OC43 and HKU1 infect through S1 binding to ubiquitous 9-O-acetyl sialic acid residues on host glycoproteins and lipids, α-CoV 229E utilizes S1 binding to aminopeptidase N (CD13), and α-CoV NL63 entry using both ACE2 and heparin sulfate proteoglycans; and β-CoV MERS-CoV uses dipeptidyl peptidase 4 (DPP4). Amongst the CoV, including β-CoV OC43 and HKU1, the precise binding sites for attachment receptors has still not been conclusively defined(26–28), suggesting possible heterogeneity or fluidity in the evolution of CoV as they adapt to humans. Although SARS-CoV-2 RBD binds with high affinity to ACE2, facilitating attachment to host cell and ultimate infection, and to date SARS-CoV-2 entry and pathology appears highly dependent on ACE2, some reports have described ACE2-independent SARS-CoV-2 infection *in vitro*. This has included acquisition of the ability for heparin sulfate mediated infection of an ACE2-negative human lung epithelial as a result of the E484D mutation in the RBD(29), indeed *in vitro* evolution of SARS-CoV-2 to higher infectivity through more efficient binding to heparin sulfate has been reported(30), and *in vivo* reduction of intestinal ACE2 expression did not impact SARS-CoV-2 infection or severity(31), further suggesting the contribution of other cellular entry mechanisms. Several other receptors have been reported to facilitate ACE2-independent SARS-CoV-2 infection, including the tyrosine-protein kinase receptor UFO (AXL)(32), LDLRAD3, and CLEC4G(33). While it is almost certain that the current SARS-CoV-2 VoC ravaging humanity are highly dependent on ACE2, as the high level of viral burden among the world’s population continues, increasingly in people with pre-existing Abs to SARS-CoV-2 through prior infection or vaccination, S2 hmAbs may provide protection against possible future variations in attachment receptor utilization by CoV.

Numerous SARS-CoV-2 RBD specific hmAbs have been approved for clinical use, and unfortunately several became irrelevant with their inability to neutralize VoC, including Omicron, highlighting the perilous future of RBD only based mAb therapeutics against CoV. The ability of 1249A8 given as a single dose to mitigate SARS-CoV-2 pathology and viral burden against all VoC tested when used either prophylactically or therapeutically, along with the added benefit of combining with a broad and potent RBD-specific hmAb substantiate its clinical potential. Further we demonstrated that direct respiratory delivery of hmAb results in potent *in vivo* activity, consistent with our and others previous finding of its more rapid and efficient distribution of mAb to the airways compared to systemic administration(16, 34). The demonstrated activity of 1249A8 against all β-CoV further highlights its clinical potential against future SARS-CoV-2 VoCs, novel β-CoVs, as well as seasonal β-CoV. With recent advances in half-life extension of clinical mAbs, formulations as stabilized dry-powder, and easy inhaled delivery; a 1249A8 and 1213H7 hmAb or similar S2 and S1 hmAb cocktail could have substantial clinical potential for β-CoV prevention and treatment. Additionally, such a prophylactic cocktail may be beneficial for immunocompromised populations that do not mount strong durable responses to current COVID-19 vaccines. The increasing availability of oral SARS-CoV-2 antivirals such as nirmatrelvir/ritonavir (paxlovid) and molnupiravir are anticipated to significantly reduce SARS-CoV-2 hospitalizations and severe infections and ultimate health care burdens of the current pandemic. Their requirements for multiple doses, short half-life, and unknown potential for occurrence of resistant viruses may limit some aspects of their clinical utility. Further, a long-acting inhaled 1249A8 and 1213H7 cocktail may be advantageous manner of passive immunization for the substantial population that remains vaccine hesitant and those that are severely immunocompromised.

Overall, we describe a potent pan-β-CoV neutralizing S2 specific human mAb, 1249A8, and demonstrated its remarkable *in vivo* activity in multiple animal models against multiple SARS-CoV-2 VoC and SARS-CoV. We further demonstrate the clear clinical potential of 12498 including in combination with a potent SARS-CoV-2 RBD-specific hmAb, 1213H7, and value direct respiratory delivery. Its occurrence indicates that universal β-CoV neutralizing hmAbs that mitigate all β-CoVs including seasonal OC43 and HKU1 can be induced and suggests further studies of such mAbs can critically inform universal CoV vaccine development.

## Materials and Methods

### Human subjects, sample collection, and B cell isolation

Peripheral blood was collected at the University of Alabama at Birmingham from adult convalescent patients approximately 1 month following PCR confirmed infection with SARS-CoV-2. The subjects provided informed consent. All procedures and methods were approved by the Institutional Review Board for Human Use at the University of Alabama at Birmingham, and all experiments were performed in accordance with relevant guidelines and regulations. Peripheral blood mononuclear cells (PBMC) were isolated by density gradient centrifugation and cryopreserved. S2-STBL and a SARS-CoV-1 / SARS-CoV-2 chimera (S1/S2) were used to generate tetramers for B-cell isolation. S2-STBL consists of the SARS-CoV-2 amino acid sequence (Wuhan-1) residues 696-1211, with mutations Q774C, L864C, S884C, A893C, K989-P, and V989P. The prefusion S1S2 chimera contains SARS-CoV-1 S residues 13-634 (uniprot P59594) and SARS-CoV-2 Wuhan-1 (P0DTC2) S residues 635-1211. The S1S2 sequence contains mutations R682S, R683-A, K989-P, and V989P. The C-termini of both proteins contain C-terminal T4 fibritin trimerization domains, his8 tags and biotinylation tags. The proteins were expressed in insect cells and purified by nickel affinity chromatography. The purified proteins were biotinylated using biotin ligase (BIRA, https://www.avidity.com/) and then used to form S2-STBL and S1S2 streptavidin tetramers for B cell isolation experiments. Cryopreserved cells were thawed and then stained for flow cytometry similar as previously described (16), using anti-CD19-APC-Cy7 (SJ25C1, BD Biosciences), HIV gp140-AlexaFluor488, S2-STBL-BV421, S1/S2 chimera-AlexaFluor647, CD3-BV510 (OKT3, Biolegend), CD4-BV510 (HI30, Biolegend), CD14-BV510 (63D3, Biolegend), CD27-PE (CLB-27/1,Life Technologies), Annexin V-PerCP-Cy5.5 (Biolegend), SA-BV421 (Biolegend), SA-AlexaFluor647 (Biolegend), and Live/Dead aqua (Molecular Probes).

### Biosafety

All *in vitro* and *in vivo* experiments with live SARS-CoV-2, SARS-CoV, and MERS-CoV were conducted in appropriate biosafety level (BSL) 3 and animal BSL3 (ABSL3) laboratories at Texas Biomedical Research Institute and Colorado State University. Experiments were approved by the Texas Biomedical Research Institute and Colorado State University Biosafety and Animal Care and Use (IACUC) committees.

### Monoclonal antibody production

Single B cells were sorted using a FACSMelody (BD Biosciences) into 96-well PCR plates containing 4 μl of lysis buffer as previously described (35). Plates were immediately frozen at −80°C after sorting until thawed for reverse transcription and nested PCR performed for IgH, Igλ, and Igκ variable gene transcripts as previously described (35, 36). Paired heavy and light chain genes were cloned into IgG1 expression vectors and were transfected into HEK293T cells and culture supernatant was concentrated using 100,000 MWCO Amicon Ultra centrifugal filters (Millipore-Sigma, Cork, Ireland), and IgG captured and eluted from Magne Protein A beads (Promega, Madison, WI) as previously described (35, 36). Immunoglobulin sequences were analyzed by IgBlast (www.ncbi.nlm.nih.gov/igblast) and IMGT/V-QUEST (http://www.imgt.org/IMGT_vquest/vquest) to determine which sequences should lead to productive immunoglobulin, to identify the germline V(D)J gene segments with the highest identity, and to scrutinize sequence properties. CV3-25, S2P6, and CC40.8 were previously described (17–19) and heavy and light chain variable regions synthesized by IDT based on reported sequences (GenBank: MW681575.1, GenBank: MW681603.1 and (19)) and cloned into IgG1 expression vector for production in HEK293T cell. 1249A8 hmAb used for *in vivo* experiments was modified to increase half-life with M252Y/S254T/T256E (YTE) mutations(37).

### Cells and Viruses

African green monkey kidney epithelial cells (Vero E6, CRL-1586) were obtained from the American Type Culture Collection. A Vero E6 cell line expressing human ACE2 and TMPRSS2 (Vero AT) was obtained from BEI Resources (NR-54970). Cells were maintained in Dulbecco’s modified Eagle medium (DMEM) supplemented with 5% (vol/vol) fetal bovine serum (FBS, VWR) and 1% penicillin–streptomycin–glutamine (PSG) solution (Corning). SARS-CoV-2 WA-1 (NR-52281), SARS-CoV-2 Beta (NR-54008), SARS-CoV-2 Gamma (NR-54982), SARS-CoV-2 Delta (NR-55611), and SARS-CoV-2 Omicron (NR-56461); SARS-CoV, Urbani strain icSARS-CoV (NR-18925); and MERS-CoV, icMERS-CoV-RFP-ΔORF5 (NR-48813) were obtained from BEI Resources. SARS-CoV-2 Epsilon was kindly provided by Dr. Charles Chiu, UCSF. The recombinant reporter-expressing SARS-CoV-2 were generated previously (21, 38).

### Binding characterization

ELISA plates (Nunc MaxiSorp; Thermo Fisher Scientific, Grand Island, NY) were coated with recombinant CoV proteins at 1□μg/ml. Recombinant proteins used include SARS-CoV-2 S2, SARS-CoV-2 S1, SARS-CoV-2 S1+S2, MERS-CoV S2, OC43 S2, HKU1 S2 (Sino Biological, Wayne, PA), and SARS-CoV S (BEI Resources). Human plasma or purified hmAbs were diluted in PBS, and binding was detected with HRP-conjugated anti-human IgG (Jackson ImmunoResearch, West Grove, PA). In select ELISAs, 8M urea were added to the ELISA plate and the plates incubated for 15□min at room temperature prior to washing with PBS plus 0.05% Tween20 and detection with anti-IgG-HRP to evaluate avidity. Immunofluorescence assay was used to determine hmAb binding to SARS-CoV-2, SARS-CoV, or MERS-CoV infected cells. Briefly, confluent monolayers of Vero E6 cells were mock infected or infected with the indicated virus. At 24□hours post infection (hpi), cells were fixed with 4% paraformaldehyde (PFA) for 30 minutes and permeabilized with 0.5% Triton X-100–PBS for 15 min at room temperature, and blocked with 2.5% Bovine Serum Albumin at 37°C for 1 h. Cells were then incubated for 1□h at 37°C with 1□μg/ml of indicated hmAb. Then, cells were incubated with fluorescein isothiocyanate (FITC)-conjugated secondary anti-human Ab (Dako) for 1□h at 37°C. Images were captured using a fluorescence microscope and camera with a 10X objective.

### Bio-layer interferometry (BLI)

Experiments for BLI were performed on a Gator Prime instrument at 30°C with shaking at 400-1000 rpm. All loading steps were 300s, followed by a 60s baseline in KB buffer (1X PBS, 0.002% Tween 20, and 0.02% BSA, pH 7.4), and then a 300s association phase and a 300s dissociation phase in K buffer. For the binding BLI experiments, mAbs were loaded at a concentration of 0.5 μg/mL in PBS onto Anti-Human IgG Fc capture (HFc) biosensors for a shift of 0.3 nm. After baseline, probes were dipped into five two-fold serial dilutions of Spike protein from SARS-CoV-2, SARS-CoV, or MERS (all from Acro Biosystems, Newark, DE) starting at 50 nM and a 0 nM for the association phase.

### Surface Plasmon Resonance (SPR)

SPR experiments were performed on a Biacore T200 (Cytiva) at 25°C using a running buffer consisting of 10mM HEPES, 150mM NaCl, 0.0075% P20. Binding comparisons between 1249A8, CC40.8, S2P6, and CV3-25 were performed by capturing the hmAbs to the chip surface of CM-5 chips using a human antibody capture kit (cytiva). The binding kinetics for the interaction between hmAbs and SARS-CoV-2 Spike protein (R&D Systems) was determined by injecting four concentrations of SARS-CoV-2 Spike (25 nM highest concentration) with a contact time of 240 seconds and a 300 second dissociation phase. The same parameters were used to characterize MERS-CoV (Sino Biologicals) binding to the hmAbs. All SPR experiments were double referenced (e.g., sensorgram data was subtracted from a control surface and from a buffer blank injection). The control surface for all experiments consisted of the capture antibody. Sensorgrams were globally fit to a 1:1 model, without a bulk index correction, using Biacore T-200 evaluation software version 1.0.

### CoV neutralization

hmAbs were tested for neutralization of live SARS-CoV-2, SARS-CoV, and MERS-CoV as previously described (39). Vero E6 cells (96-well plate format, 4 × 10^4^ cells/well, quadruplicate) were infected with 100-200 PFU/well of SARS-CoV-2. SARS-CoV-2 Omicron neutralization was performed in Vero AT using 600 PFU/well. After 1 h of viral adsorption, the infection media was changed with the 100 μl of postinfection media containing 1% Avicel and 2-fold dilutions, starting at 25 μg/ml of hmAb (or 1:100 dilution for human serum control). At 24 h p.i., infected cells were fixed with 10% neutral formalin for 24 h and were immune-stained using the anti-NP monoclonal antibody 1C7C7 (39). Virus neutralization was evaluated and quantified using ELISPOT, and the percentage of infectivity calculated using sigmoidal dose response curves. The formula to calculate percent viral infection for each concentration is given as [(Average # of plaques from each treated wells – average # of plaques from “no virus” wells)/(average # of plaques from “virus only” wells - average # of plaques from “no virus” wells)] x 100. A non-linear regression curve fit analysis over the dilution curve can be performed using GraphPad Prism to calculate NT_50_. Mock-infected cells and viruses in the absence of hmAb were used as internal controls. hmAbs were also tested using a SARS-CoV-2 Spike protein pseudotyped virus (PsV) expressing firefly luciferase. Virus neutralization was measured by the reduction of luciferase expression. VeroE6/TMPRSS2 cells were seeded at 2 × 10^4^ cells/well in opaque plates (Greiner 655083). The next day, PsV corresponding to 1-10 × 10^6^ luciferase units was mixed in Opti-MEM with dilutions of hmAbs and incubated at RT for 1 h. Media was removed from the cells and 100 μl/well of the hmAb/PsV mix was added in triplicates. After 1 h incubation at 37C and 5% CO_2_, another 100 μL of Opti-MEM was added, and cells were incubated for 24 more hours. After this time, luciferase activity was measured using Passive Lysis Buffer (Promega E1941) and Luciferase substrate (Promega E151A) following the manufacturer’s instructions. Neutralization was calculated as the percent reduction of luciferase readings as compared to no-antibody-controls.

### Antibody-Dependent Cellular Phagocytosis (ADCP) assay

ADCP activity of the mAbs was measured as previously described (*36, 40*) with slight modifications. Briefly, SARS-CoV-2 Wuhan-Hu-1 Spike protein (NR-53524 BEI Resources) or MERS-CoV Spike protein (Sino Biological) was biotinylated with the Biotin-XX Microscale Protein Labeling Kit (Life Technologies, NY, USA). 0.25 μg of biotinylated Ag or ~0.16 μg of BSA (used as a baseline control in an equivalent number of Ag molecules / bead) was incubated overnight at 4°C with 1.8 × 10^6^ Yellow-Green neutravidin-fluorescent beads (Life Technologies) per reaction in a 25 μL of final volume. Antigen-coated beads were subsequently washed twice in PBS-BSA (0.1%) and transferred to a 5 mL Falcon round bottom tube (Thermo Fisher Scientific, NY, USA). mAbs, diluted at 5 μg/ml, were added to each tube in a 20 μL of reaction volume and incubated for a 2 h at 37°C in order to allow Ag-Ab binding. Then 250,000 THP-1 cells (human monocytic cell line obtained from NIH AIDS Reagent Program) were added to the cells and incubated for 3 h at 37°C. At the end of incubation, 100 μL 4% paraformaldehyde was added to fix the samples. Cells were then assayed for fluorescent bead uptake by flow cytometry using a BD Biosciences Symphony. The phagocytic score of each sample was calculated by multiplying the percentage of bead positive cells (frequency) by the degree of phagocytosis measured as mean fluorescence intensity (MFI) and dividing by 10^6^. Values were normalized to background values (cells and beads without mAb) and an isotype control to ensure consistency in values obtained on different assays. Finally, the phagocytic score of the testing mAb was expressed as the fold increase over BSA-coated beads.

### K18 hACE2 transgenic mice experiments

All animal protocols involving K18 hACE2 transgenic mice were approved by the Texas Biomedical Research Institute IACUC (1718MU). Five-week-old female K18 hACE2 transgenic mice were purchased from The Jackson Laboratory and maintained in the animal facility at Texas Biomedical Research Institute under specific pathogen-free conditions and ABSL3 containment. For virus infection, mice were anesthetized following gaseous sedation in an isoflurane chamber and inoculated with viral dose of 10^5^ PFU per mouse, intranasally. For *ex vivo* imaging of lungs, mice were humanely euthanized at 2 and 4 d p.i. to collect lungs. Fluorescent images of lungs were photographed using an IVIS (AMI HTX), and the brightfield images of lungs were taken using an iPhone 6s (Apple). Nasal turbinate and lungs from mock or infected animals were homogenized in 1 mL of PBS for 20 s at 7,000 rpm using a Precellys tissue homogenizer (Bertin Instruments). Tissue homogenates were centrifuged at 12,000 × g (4 °C) for 5 min, and supernatants were collected and titrated by plaque assay and immunostaining as previously described. For the body weight and survival studies, five-week-old female K18 hACE2 transgenic mice were infected intranasally with 10^5^ PFU per animal following gaseous sedation in an isoflurane chamber. After infection, mice were monitored daily for morbidity (body weight) and mortality (survival rate) for 11 d. Mice showing a loss of more than 25% of their initial body weight were defined as reaching the experimental end point and humanely euthanized.

### Golden Syrian hamster experiments

Experiments using Syrian hamsters were approved for use by the Colorado State University IACUC. For each experiment, forty female Syrian hamsters (Envigo Corporation, Indianapolis, IN, USA) were housed in ventilated cages in separate rooms under ABSL3 containment. They were acclimated for 7-14 days after arrival and inoculated with virus at 6 weeks of age. Animals were challenged with virus under ketamine-xylazine anesthesia by intranasal instillation of 100 ul of virus diluted in PBS to achieve a dose of 10^4^ CCID50 ((rSARS-CoV-2 Delta B.1.617.2 (hCoV-19/USA/CA-VRLC086/2021) BEI Resources (Manassas, VA, USA) or plaque-forming units (SARS-CoV, Urbani Strain; BEI Resources); the inocula were back-titrated after completion of the challenge to confirm dose delivered. Antibody therapies were administered as summarized in Figure 7 and 8 at 12 hours following virus inoculation, again under ketamine-xylazine anesthesia and by intranasal instillation of 100 ul. Oropharyngeal swabs were collected daily from all hamsters on days 1, 2 and 3 post-virus inoculation. Swabs were broken off into 1 ml of BA1 medium (Tris-buffered minimal essential medium containing 1% BSA) supplemented with 5% fetal bovine serum (BA1-FBS) and stored at −80C until assay. Half of the hamsters inoculated with virus were euthanized on day 3 and half on day 7 post-challenge. For the animals euthanized on day 3, samples of nasal turbinates and cranial and caudal right lung were homogenized in BA1-FBS using a mixer mill and stainless-steel balls to obtain ~10% tissue homogenates. Infectious virus in tissue homogenates and oropharyngeal swabs was titrated by double-overlay plaque assay. Briefly, 10-fold serial dilutions of samples were prepared in BA1 medium with antibiotics, inoculated onto confluent monolayers of Vero cells in 6-well plates, incubated with rocking for 45 minutes, and then overlaid with 0.5% agarose in phenol-red free MEM supplemented with antibiotics. Plates were incubated for one (SARS CoV) or two (SARS-CoV-2) days and second overlay containing neutral red dye was added. Plaques were counted 1 day after the second overlay. The limit of detection for this assay was 10 PFU/swab and 100 PFU/gram of tissue.

### Statistical analysis

Significance was determined using GraphPad Prism, v8.0. Twotailed t-test were applied for evaluation of the results between treatments. p<0.05 was considered significant. For statistical analysis viral titers were log transformed and undetectable virus was set to the limit of detection.

## Acknowledgments

We are grateful for the clinical research staff that enabled this project and for the technical assistance provided by Christopher Bates, the technical guidance provided by Justin Roth, the virology expertise provided by Ilya Frolov, and the assistance of the University of Alabama at Birmingham (UAB) Center for AIDS Research and its Flow Cytometry Core Facility under the direction of Olaf Kutsch and the UAB Multidisciplinary Molecular Interaction Core Facility under the direction of Randall Davis. We thank Alexander Rosenberg and Christopher Fucile for providing immunoglobulin sequence analysis software. We thank Rachel Maison and Airn Hartwig for valuable assistance in conducting the hamster experiments. We are most grateful for the participation of the study volunteers. Funding for this work provided by institutional support from the University of Alabama at Birmingham (to J.J.K.) and the Texas Biomedical Research Institute (to L.M.-S.), funding from Aridis Pharmaceuticals, and the National Institutes of Health (1R01AI161175 to J.J.K., L-M.S, and M.R.W.) and funded Center for Research on Influenza Pathogenesis and Transmission (CRIPT), a NIAID-funded Center of Excellence for Influenza Research and Response (CEIRR, contract # 75N93021C00014 to L.M-S.), and NIAID Preclinical and Clinical Services Contracts Division of Microbiology and Infectious Diseases, Respiratory Diseases Branch.

## Data Availability

All relevant data are within the manuscript.

## Author contributions

J.J.K., L.M.-S., M.R.W., R.A.B., and V.L.T. conceived the study. M.S.P., M.B., S.S., and J.J.K. isolated and screened the mAbs. M.S.P., J.J.K., M.R.W., and A.D. produced the mAbs. M.R.W. and A.D. designed and produced the recombinant S2 proteins and conducted the SPR analysis. M.B. conducted ADCP. J.-G.P., C.Y., R.A.B. and L.M.-S. conducted the *in vitro* and *in vivo* antiviral assessments. I.F. conducted *in vitro* antiviral assessments. R.A. B. conducted *in vivo* antiviral assessments. P.A.G. and N.B.E. acquired the clinical specimens. A.L. conducted the pseudovirus neutralization assay and BLI. J.W., P.L., and V.L.T. designed and conducted the experiments to determine the efficacy of mAbs. D.C., J.W., and P.L. provided expert advice. J.J.K., L.M.-S., M.R.W., R.A.B., and V.L.T. supervised the work. J.J.K., L.M.-S., M.R.W., R.A. B., and V.L.T. wrote the manuscript. All of the authors reviewed the manuscript.

